# Design principles of selective transport through biopolymer barriers

**DOI:** 10.1101/709675

**Authors:** Laura Maguire, Michael Stefferson, Meredith D. Betterton, Loren E. Hough

## Abstract

In biological systems, polymeric materials block the movement of some macromolecules while allowing the selective passage of others. In some cases, binding enables selective transport, while in others the most inert particles appear to transit most rapidly. To study the general principles of filtering, we develop a model motivated by features of the nuclear pore complex (NPC) which are highly conserved and could potentially be applied to other biological systems. The NPC allows selective transport of proteins called transport factors which transiently bind to disordered, flexible proteins called FG Nups. While the NPC is tuned for transport factors and their cargo, we show that a single feature is sufficient for selective transport: the bound-state motion resulting from transient binding to flexible filaments. Interchain transfer without unbinding can further improve selectivity, especially for crosslinked chains. We generalize this observation to model nanoparticle transport through mucus and show that bound-state motion accelerates transport of transient nanoparticle application, even with clearance by mucus flow. Our model provides a framework to control binding-induced selective transport in bipolymeric materials.

## INTRODUCTION

Living systems control the localization and movement of molecules, nanoparticles, viruses, and other organisms using selective filters made of biopolymers [1]. These filters regulate access to genetic material (the nuclear pore complex, or NPC), cells (the pericellular matrix), tissues (the extracellular matrix), and organs (mucus). In addition to their protective role, polymeric biomaterials can inhibit delivery of therapeutic agents. How particle binding affects motion and filtering is unclear: binding to the pericellular matrix facilitates uptake of nanoparticles by single cells, and transport factors that bind to proteins in the NPC move rapidly through it [2, 3]. In contrast, binding inhibits the uptake of nanoparticles that bind to airway mucus, and many viruses minimize binding interactions [4–8]. Mucus presents a particularly formidable challenge to nanoparticle drug delivery, where vectors tend to be of a similar size to pathogens, the exclusion of which is the primary role of mucus. While particle size, charge, and binding interactions are known to affect filtering, the physical principles that underlie mobility and transport in polymeric biomaterials are not fully understood [1].

The nuclear pore complex (NPC) relies on binding for selective transport and is important for diverse cellular processes including gene regulation and translation [3]. The NPC selectively filters molecular traffic between the nucleus and cytoplasm of eukaryotic cells, preventing the passage of most macromolecules while allowing the rapid passage of others of similar size and charge. A defining feature of molecules allowed passage is that they bind, either directly or indirectly, to the selective barrier of the NPC. Transport occurs through the central channel, ~50 nm in diameter and ~100 nm long. The selective barrier filling the central channel is made from disordered proteins, the FG nucleoporins (FG Nups), which contain repeated phenylalanine-glycine (FG) motifs (Fig. 1). Transport factors that directly bind to the FG repeats can cross the NPC and carry cargo with them [3]. Transport through the NPC is remarkably fast, with pore residence times ~10 ms [9]. Consistent with this, the binding kinetics of FG Nup-transport factor interactions are rapid, often approaching diffusion-limited [10, 11].

**Figure 1.**
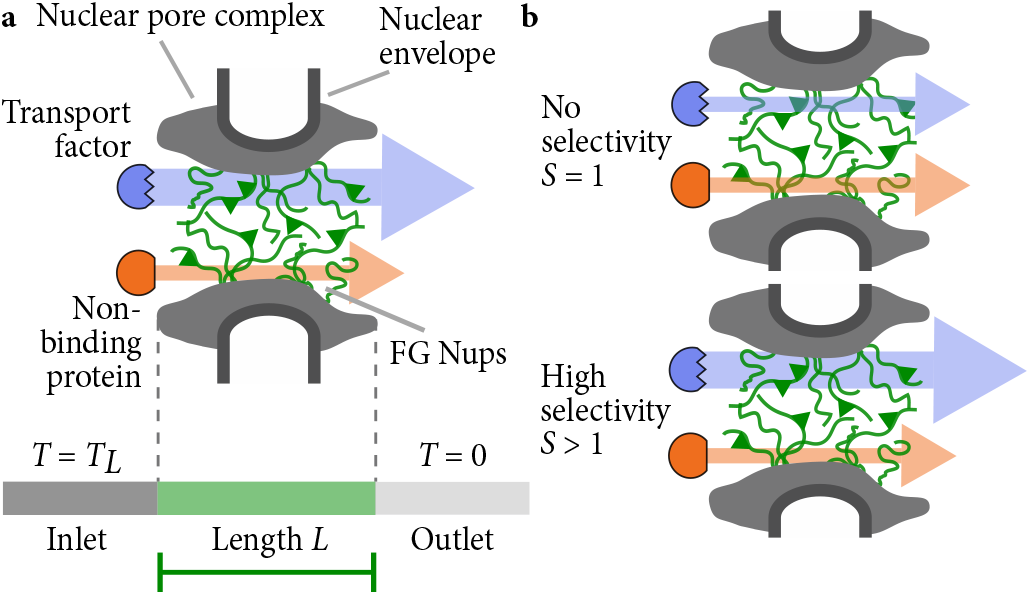
Schematics of the nuclear pore complex and model. (a) The nuclear pore complex (grey) is filled with FG Nups (green polymers) that selectively passage transport factors that bind to FG Nups (blue) while blocking non-binding proteins (red). The central channel of the pore has length *L*. Protein concentration is high on the left (inlet) and low on the right (outlet). (b) Selectivity quantifies the degree of selective transport through the pore. A non-selective pore with *S* = 1 has the same flux for a transport factor as for a non-binding protein (top). A selective pore with *S* > 1 has a larger flux for a transport factor than a non-binding protein (lower).

Current optimally-designed nanoparticles for drug delivery through mucus minimize binding interactions, a striking difference from nuclear transport. These mucus-penetrating particles, with minimal interactions with mucus, have enhanced delivery properties over those that bind to lung, eye, and vaginal mucosa [4, 6, 12, 13]. In contrast, the flagella or pili of a number of bacteria bind to mucus, suggesting a possible role for binding interactions in helping pathogens pass through mucus barriers [14–17]. This raises the question of which features of binding contribute to the passage of bacteria through mucus barriers and whether those mechanisms could be applied to nanoparticles as well.

Binding does not always enhance flux across a barrier. Mechanisms of selective transport include transport by mobile carriers and particle motion directly from one binding site to the next [18–23]. In general, selectivity requires that particles move while bound. Systems without binding, such as oil barriers separating aqueous reservoirs, have higher flux as a consequence of higher concentration within the barrier (Fig. S2 [24]). Binding-based concentration is the basis of several models of selective transport in the NPC which represent binding and diffusion using an effective free-energy landscape of the barrier to passage. In this picture, binding of transport factors to FG Nups provides a free-energy gain which offsets the energy barrier and is presumed to allow for more effective transport [25]. Details of the free energy landscape have seen extensive exploration both *in vivo* and *in vitro* [26–30], but previous work has not typically investigated the underlying molecular mechanisms.

In this paper we study molecular mechanisms arising from simple, conserved features of the NPC that could potentially be applied to other biological and synthetic systems. Biomimetic filters or drug-delivery vehicles lack the full complexity of the FG barrier or FG Nup-transport factor interactions. Therefore, we focus on two features of binding between some transport factors and FG repeats: First, binding is diffusion-limited and transient between individual FG repeats and transport factors [10, 11]. While some transport factors bind more tightly, such transport factor-FG interactions typically require active release [31]. Second, FG Nups remain fully disordered and highly dynamic upon binding [10, 11, 32, 33]. We developed a model to determine whether these two features are sufficient to produce binding-induced selective transport.

To compare the contributions of different molecular mechanisms of selective transport, we develop a hierarchy of models, beginning with a continuum model of bound-state diffusion. This model confirms that bound mobility is sufficient for selective transport. Then we consider two molecular mechanisms of bound diffusion: first a model of diffusion with transient binding to flexible polymers, and second a model of multivalent binding interactions that allow inter-chain transfers while bound. Finally, we extend the continuum model to include features specific to mucus barriers, including fluid clearance and mucus sloughing to study how these effects modify selectivity arising from bound-state diffusion.

## BOUND MOBILITY CONTROLS SELECTIVITY IN A MINIMAL MODEL OF TRANSPORT THROUGH A POLYMER BARRIER

To understand how binding can cause selective transport, we formulated a minimal model. Because we are using the NPC as our model system, we use NPC-related terminology. However, to keep our model relevant to a wider range of biopolymer filters, we neglect NPC-specific features and consider only the effect of multivalent binding to flexible tethers. The NPC-specific features we neglect include a wide capture area [34], varying landscape along the axis of the NPC [26–29], varying pore composition [35–38], active release from the pore [28, 39, 40], and modulation of FG Nup dynamics by transport factors [41, 42].

To compare model predictions to experiment, we consider the prototypical transport factor nuclear transport factor 2 (NTF2), a homodimer that selectively passes through the nuclear pore without active release. It is the smallest known transport factor. While similarly-sized nonspecific proteins can passage the NPC, the inert transport rate is significantly reduced: NTF2 is estimated to have 30-120 times higher flux through the NPC than GFP [43–45]. NTF2 may be less affected by transient crosslinking with the NPC because of its small size.

Although it is typically assumed that transport factors within the NPC are always bound to FG Nups, binding is sufficiently rapid that there can be many unbinding events during transport. The dissociation constant of NTF2 for FG Nups has been measured at values between nanomolar and millimolar [11, 46–50]. Assuming that binding is diffusion limited, the slower end of this range corresponds to an unbinding rate of 1 s^−1^, incommensurate with transport in the absence of active release. However, the fastest unbinding rate is 10^−6^ s^−1^, allowing for up to 10^4^ unbinding events during a 10-ms transport event. If the measured K_*D*_ recently obtained for the interaction of NTF2 with a specific portion of the FG Nup Nsp1 (1 mM) [50], is representative of all NTF2-FG interactions, then NTF2 is expected to spend approximately 50% of its time unbound while in the NPC.

We model transport factor diffusion within the NPC including binding kinetics and diffusion of the transport factor in both free and bound states. We focus on transport within the pore, since in single-molecule measurements most of the transport time is spent in a random walk within the central channel [9, 29]. Here we consider the limit in which binding kinetics provide the only barriers to entry and exit from the pore. (See Supplemental Material [24] for treatment of rate-limiting entry and exit.) We impose a fixed transport factor concentration difference across the barrier, consistent with passive transport. In mucosal and biomimetic systems, the exterior concentration would be imposed by the delivery mechanism, which is well described by this model.

In order to determine the degree of selective transport resulting from bound diffusion, we consider a barrier of length *L* that separates two reservoirs [Fig. 1(a)]. Within the barrier are free transport factor (concentration *T*), free FG Nups (*N*), and bound transport factor-FG Nup complex (*C*), with total FG Nup concentration *N_t_* = *N* + *C*. Transport factor diffusion within the barrier (0 < *x* < *L*) is described by the reaction-diffusion equations

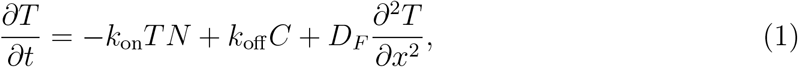

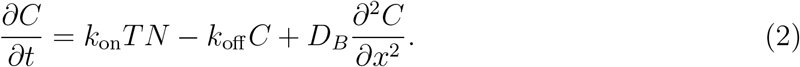

Transport factor-FG Nup binding has on-rate constant *k*_on_, off-rate *k*_off_, and dissociation constant *K_D_* = *k*_off_/*k*_on_. We include competition between transport factors for FG binding sites and assume that the barrier properties are independent of transport-factor concentration [41, 51, 52]. The diffusion constants of free (*D_F_*) and bound (*D_B_*) transport factors are spatially constant. The fixed reservoir transport factor concentrations are *T_L_* (inlet, left) and 0 (outlet, right).

We numerically integrated the full nonlinear equations to find the flux of transport factor across the barrier, *J* = −*D_F_ ∂T/∂x*|_*x=L*_ [24]. Because flux measured in experiments is typically linearly proportional to transport factor concentration, transport factors likely remain below binding saturation in the NPC [53, 54]. Therefore, we also solved Eqs. (1, 2) analytically in the low-binding limit [24]. We define the transport selectivity *S* as the ratio of the steady-state flux of a binding versus an otherwise identical non-binding species [Fig. 1(b)]

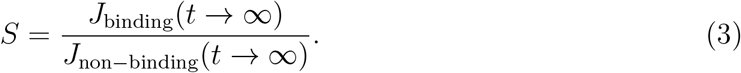

Two key differences distinguish our model from that of Yang *et al*., who also explicitly considered bound diffusion [21]. First, we solve the full nonlinear equations. Second, we enforce the requirement that transport factors unbind before leaving the pore, which has significant effects. Omitting this restriction overestimates selectivity for rapid bound diffusion and underestimates it for slow bound diffusion (Figs. S1, S12 [24]).

Our model shows that selective transport is not possible either at steady-state or during initial transient dynamics if bound transport factors do not move (*D_B_* = 0). If *D_B_* = 0, the steady-state flux *J* = *D_F_T_L_*/*L* for both binding and non-binding proteins, so *S* =1 (Fig. 2, Fig. S1 [24]). While the binding transport factor accumulates within the pore, its immobility means transport is not increased compared to the non-binding case. In other words, the concentration within the barrier is increased by binding, but the mobility and therefore flux of particles is proportionately reduced [55]. Notably, this effect is independent of binding kinetics. Prior to steady state, binding slows transport [Fig. 2(a)]. In systems such as airway mucus, immobilization by binding may increase the time available for degradation or active clearance, consistent with the observation that binding tends to inhibit selective transport in those systems [4–6]. This effect is related to the binding-site barrier seen in antibody delivery to tumors [56], and observations that non-binding nanoparticles are often more effective in drug delivery to tumors than binding particles [1].

**Figure 2.**
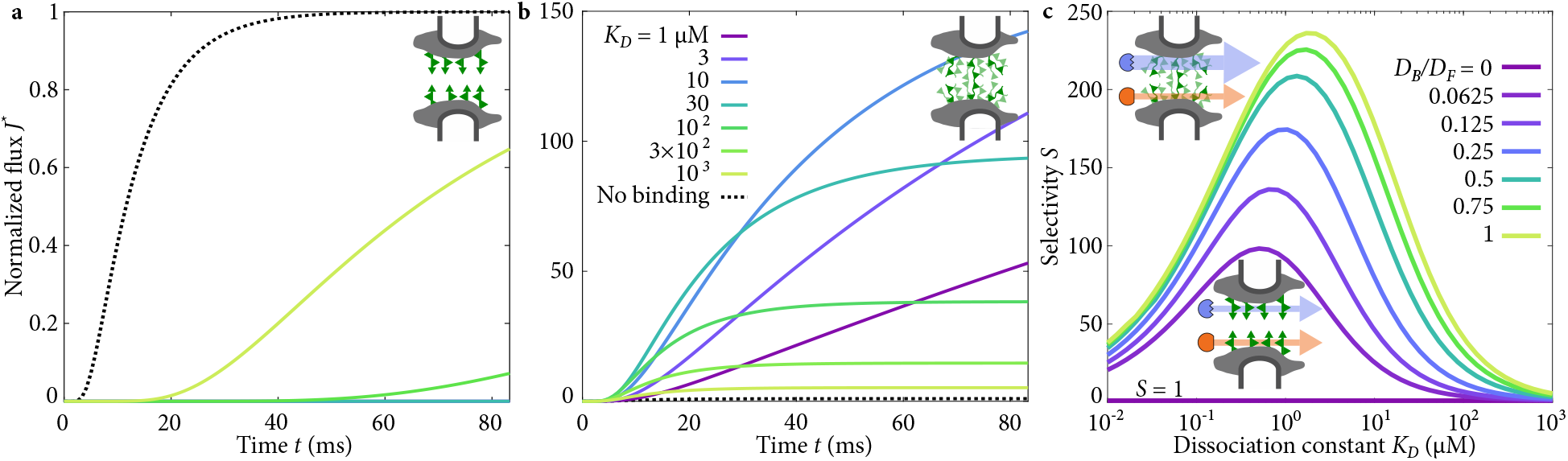
Flux through the pore and selectivity for transport factors with varying bound mobility. (a) Flux as a function of time when transport factors are immobile while bound, with varying binding affinity as in (b). (b) Flux as a function of time when transport factors are mobile while bound with *D_B_* = *D_F_*, with varying binding affinity. Note change in y-axis scale. (c) Selectivity as a function of dissociation constant with varying bound diffusion coefficient.

When bound transport factors are mobile (*D_B_* > 0), transport is enhanced for the binding particle: it is selective by a factor of up to 240 for experimentally relevant parameters [Fig. 2(b,c) and Supplemental Material Fig. S1 [24]]. We examined how binding kinetics influence transport and found an optimal dissociation constant of *K_D_* ~ 1 *μ*M for high selectivity [Fig. 2(c)]. We allow *K_D_* to vary, given that the affinity of NTF2 for relatively few FG Nups has been reproducibly measured [11, 47, 48, 50]. Selectivity decreases for high *K_D_* because binding is too weak to significantly increase transport factor concentration in the pore. For low *K_D_*, tight binding causes the concentration of bound complexes to become approximately constant across the pore. Because diffusive flux is driven by a concentration gradient, washing out the gradient by tight binding decreases flux and selectivity.

## BOUND DIFFUSION DUE TO TRANSIENT BINDING TO DYNAMIC POLYMERS

We next examined molecular mechanisms in bound diffusion, beginning with a model incorporating rapid binding to flexible polymers. Previous measurements have found that FG Nups are flexible and dynamic [10, 33, 57], which would allow a transport factor bound far from the anchored end of an FG Nup to move while bound. The bound motion depends on the polymer properties and binding kinetics, with more flexible polymers and more transient binding resulting in higher bound diffusion.

To quantify mobility while bound to flexible polymers, we developed a minimal model. Flexible polymers behave as entropic springs if they are not highly stretched [58]. Therefore, a bound transport factor that diffuses while attached to a spring-like tether can be represented as diffusion in a harmonic potential well (Fig. 3). The width of the harmonic well is related to the effective length of the flexible domain: if FG Nups are not crosslinked, the effective length is the full FG Nup length, while if they are crosslinked or entangled, the length is reduced [43]. The probability density of the position of a transport factor that binds to a well with center *x* = 0 is 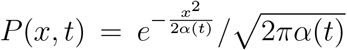, where *α*(*t*) = (1 − *e*^−2*kD_F_βt*^)/(*kβ*), *k* is the spring constant of FG Nup tethering and *β* = 1/*k_B_T* is the inverse thermal energy [59]. The transport factor mean-squared displacement (MSD) is then 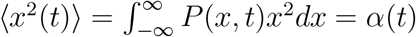. The typical MSD during a binding event depends on the probability density of binding lifetime *ρ*(*t*) = exp(−*t*/*τ*)/*τ*, where *τ* = 1/*k*_off_ is the mean bound lifetime:

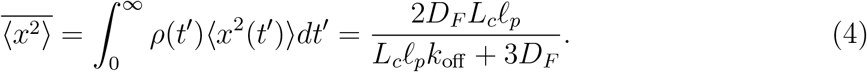

**Figure 3.**
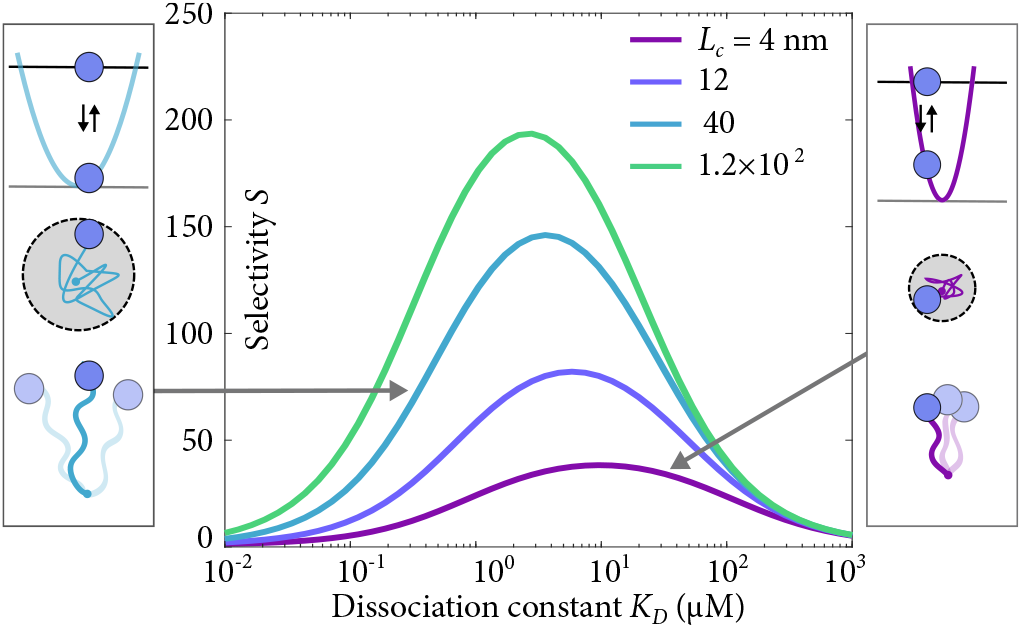
Selectivity as a function of dissociation constant, with varying polymer length in the tethered-diffusion model. Schematics of FG Nups that constrain the motion of transport factors less (left, longer FG Nup) or more (right, shorter FG Nup), which corresponds to changing the width of the harmonic potential.

Here we assume that the spring constant is that of a worm-like chain polymer *k* = 3/(2*βL_c_ℓ_p_*), where *L_c_* is the contour length and *ℓ_p_* the persistence length [58].

Because transport factor-FG Nup interactions have fast binding kinetics [10, 11], we estimate the bound diffusion coefficient by averaging over many binding events, while considering only bound motion, giving [24]

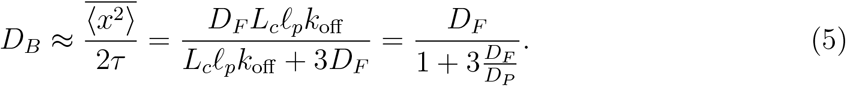

Here *D_P_* = *L_c_ℓ_p_k*_off_ quantifies how polymer properties tune the bound-state diffusion coefficient: bound mobility increases with increasing chain length or persistence length, or decreasing binding lifetime (Figs. 3, S5, S6). When *D_P_* is large (*D_F_*/*D_P_* ≪ 1), *D_B_* approaches *D_F_*, since the long chains barely affect transport factor motion during the short binding event. For small *D_P_*(*D_F_*/*D_P_* ≫ 1), transport factor motion is inhibited by a short tether, giving *D_B_* ≈ *D_P_*/3 ≪ *D_F_*. This shows that the kinetics of transport factor-FG Nup interaction are a primary determinant of the bound mobility: the faster the binding kinetics, the higher the bound diffusion constant. These relationships between microscopic binding properties and selectivity are possible because we explicitly consider the process of binding, in contrast to previous work [28].

A strength of this simple model is that we can quantitatively predict the bound diffusion constant based on polymer properties and binding kinetics. Flexible disordered proteins typically have low persistence length, *ℓ_p_* ≈ 1 nm [60]. If the on-rate constant is diffusion-limited, *k*_on_ = 10^−3^*μ*M^−1^ *μ*s^−1^ [10, 11], then the binding affinity determines the off rate. Disordered FG Nups have *L_c_* ≈ 100-280 nm, (250-700 amino acids in length [61] with a contour length of approximately 0.4 nm per amino acid). For fixed *D_F_*, we predict that selectivity is increased by increasing the binding on-rate constant *k*_on_ (Fig. S3). Thus rapid kinetics increase both the effective bound diffusion coefficient and the selectivity. Large *k*_on_ makes transport more selective because fast binding kinetics relative to diffusive motion are necessary to maintain a steep concentration gradient in the pore. High FG Nup concentration (as measured experimentally) leads to large *N_t_* and low *D_F_*, both of which increase selectivity. Decreasing *D_F_* or increasing the length of the pore both reduce the magnitude of the flux and increase selectivity (Figs. S3, S4). Therefore, varying transport factor free diffusion coefficient and pore length involves a trade-off between transit time and selectivity.

Our model provides a quantitative tool to evaluate selective transport within NPC mimics made directly from FG Nups. Because binding does not always lead to selective transport, we analyze whether these synthetic materials would act as selective filters. Materials formed *in vitro* by spontaneous self-assembly of FG Nups [62] or transient crosslinking by alpha-helical peptides [63] show strong selective *entry*. Using published data, we used our theory to predict whether these gels would show selective *transport* through a 100-nm barrier (Table S1). Predicted selectivity is ~ 10, with the predicted selectivity of one hydrogel *S* ≈ 200, apparently the most selective synthetic gel to date [62].

Using a conservative set of parameters, our tethered diffusion model predicts values of selectivity which agree with experimental measurements in the NPC (Table I). Our results explain how key features of some transport factor-FG interactions result in selective flux through the NPC. Bound transport factors are mobile due to rapid binding kinetics and thermally-driven diffusion while bound to flexible polymers. These results suggest that features such as active release, dissolution of crosslinks within a gel phase, and a wide capture area are not necessary for selective transport of NTF2 [34, 39, 62].

**Table I.** Comparison between experimental results for NTF2 and GFP and model predictions. Flux measured in molecules per pore per second.

| Method,<br>Cell type | Species | Flux | Selectivity | Notes |
| --- | --- | --- | --- | --- |
| OSTR,<br>*Xenopus* | NTF2 | 91–123 | 24–37 | [44] |
|  | GFP | 3.3–3.8 |  |  |
| OSTR,<br>*Xenopus* | NTF2 | 47.3 | 43 | [45] |
|  | GFP | 1.1 |  |  |
| Permeabilized<br>cells, HeLa | NTF2 | 250 | 125 | [43] |
|  | GFP | 2 |  |  |
| Model | Binding | 2–480 | 1–240 | This<br>work |
|  | Non-binding | 2 |  |  |

## BOUND DIFFUSION DUE TO INTER-CHAIN TRANSFER

In addition to tethered diffusion, bound motion can arise from multivalent FG Nup-transport factor interactions that allow transfer between polymer chains while remaining bound [41, 64]. The multivalent nature of transport factor-FG interactions has been assumed to be a key contributor to selectivity because a transport factor can bind simultaneously to more than one FG Nup, moving hand-over-hand while remaining bound [22, 29, 42, 48]. Consistent with this, transport factors may slide between nearby FG sites rather than fully unbinding and re-binding [22]. If the newly-bound FG repeat is on a neighboring chain, the FG tether site that constrains transport-factor motion moves while the transport factor remains bound.

To determine the relative contributions to selectivity of inter-chain transfer and tethered diffusion, we modeled both mechanisms. We simulated a transport factor that undergoes tethered diffusion when bound to an FG Nup and can directly transition between neighboring, randomly distributed tethers without unbinding (Fig. 4, [24]). Intra-chain transfers do not change the flux; therefore we neglect them.

**Figure 4.**
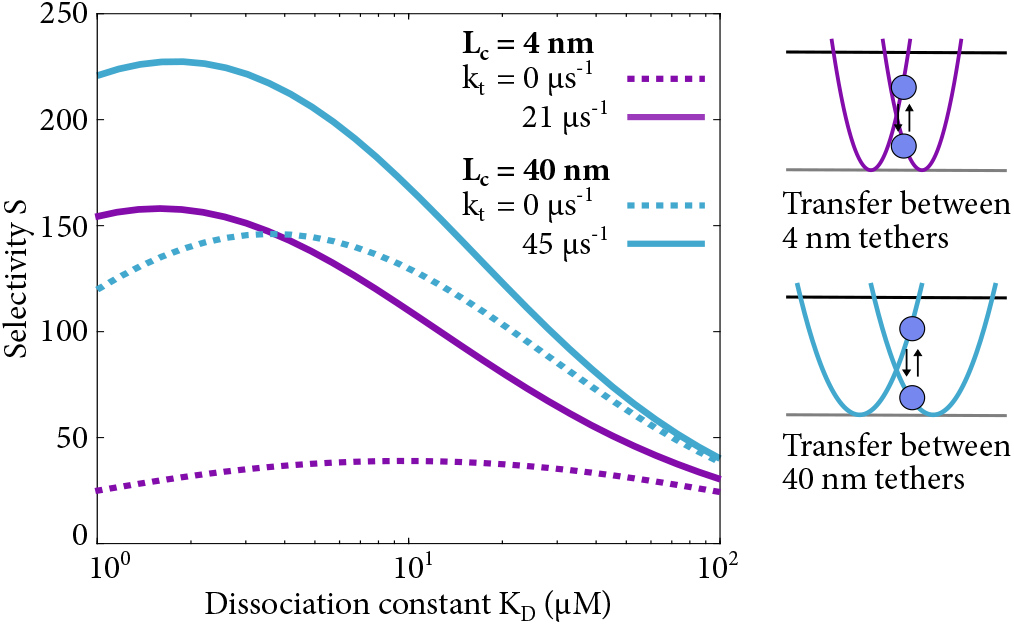
Selectivity as a function of *K_D_* with and without inter-chain transfer for FG Nup contour lengths *L_c_* = 4 nm and *L_c_* = 40 nm. FG Nups are entropic springs that constrain the motion of transport factors, and inter-chain transfer allows a transport factors to move from one FG Nup to another without unbinding at rate *k_t_*, which corresponds to switching from one harmonic well to another.

Inter-chain transfer increases selectivity by up to a factor of 6 for tight binding and short chains, which is the regime where tethered diffusion provides only limited selectivity (Figs. 4, S8, S9). For weaker binding and longer chains, inter-chain transfer leads to a modest increase in selectivity of up to 30%.

## EFFECTS OF BARRIER AND RESERVOIR DYNAMICS ON TRANSPORT IN MUCOSAL SYSTEMS

Bound diffusion might be relevant to other biopolymer filters such as mucus barriers. In contrast to the NPC, current optimally-designed nanoparticles for delivery through mucus lack binding interactions [4, 8, 13, 65]. Those particles that do bind are less effective at delivery to the underlying tissue. However, these particles were not designed with the explicit aim of bound diffusion: previously-designed particles either contained charged surfaces to interact non-specifically with highly charged mucus or contained monovalent binding motifs which would likely become immobilized while bound. As a result of these binding interactions, such particles are typically observed at the mucosal surface and are unable to penetrate. For antibody-based delivery of pharmacological agents this restriction has been termed the binding-site barrier and may explain the efficacy of antiviral antibodies whose weak mucus binding is increased upon multivalent antigen interactions [56, 66].

Our model suggests that nanoparticles could be designed to take advantage of bound diffusion in order to more effectively penetrate mucus barriers. For example, large numbers of very weakly binding motifs could coat nanoparticle-based delivery vehicles, or those motifs could be attached to nanoparticles via flexible polymers.

Mucus dynamics differ from the NPC because mucus and the surrounding fluid are cleared, potentially affecting the efficacy of selective transport arising from bound diffusion. Therefore we consider two additional features in our model. First, liquid on the surface of the mucus is often only present transiently. For example, our tears and blinking clear the liquid from the surface of the tear film in the eye [68]. Second, even for mucosal surfaces which enable more long-term exposure to external agents, the mucus grows and is shed [69].

### Transient drug application

To model the effect of clearance of the surrounding fluid on transport kinetics, we extend our model to study the transient application of a concentration gradient. A substance (T) is introduced to the mucosal surface (N) for a short time *t_d_*, then is removed. For *t_d_* = 2 s, a constant nonzero concentration of drug is applied to the exterior of the mucosal layer (the fluid in Fig. 5). For *t* > *t_d_*, that liquid is cleared, and the concentration of drug is returned to zero, mimicking the clearance of the surrounding fluid, *e.g*., due to blinking. To focus on clearance, we retain the approximation that the mucus is both homogeneous and stationary, and the cells are assumed to be perfectly absorbing. We selected parameters consistent with those in the literature for the eye tear film [24].

**Figure 5.**
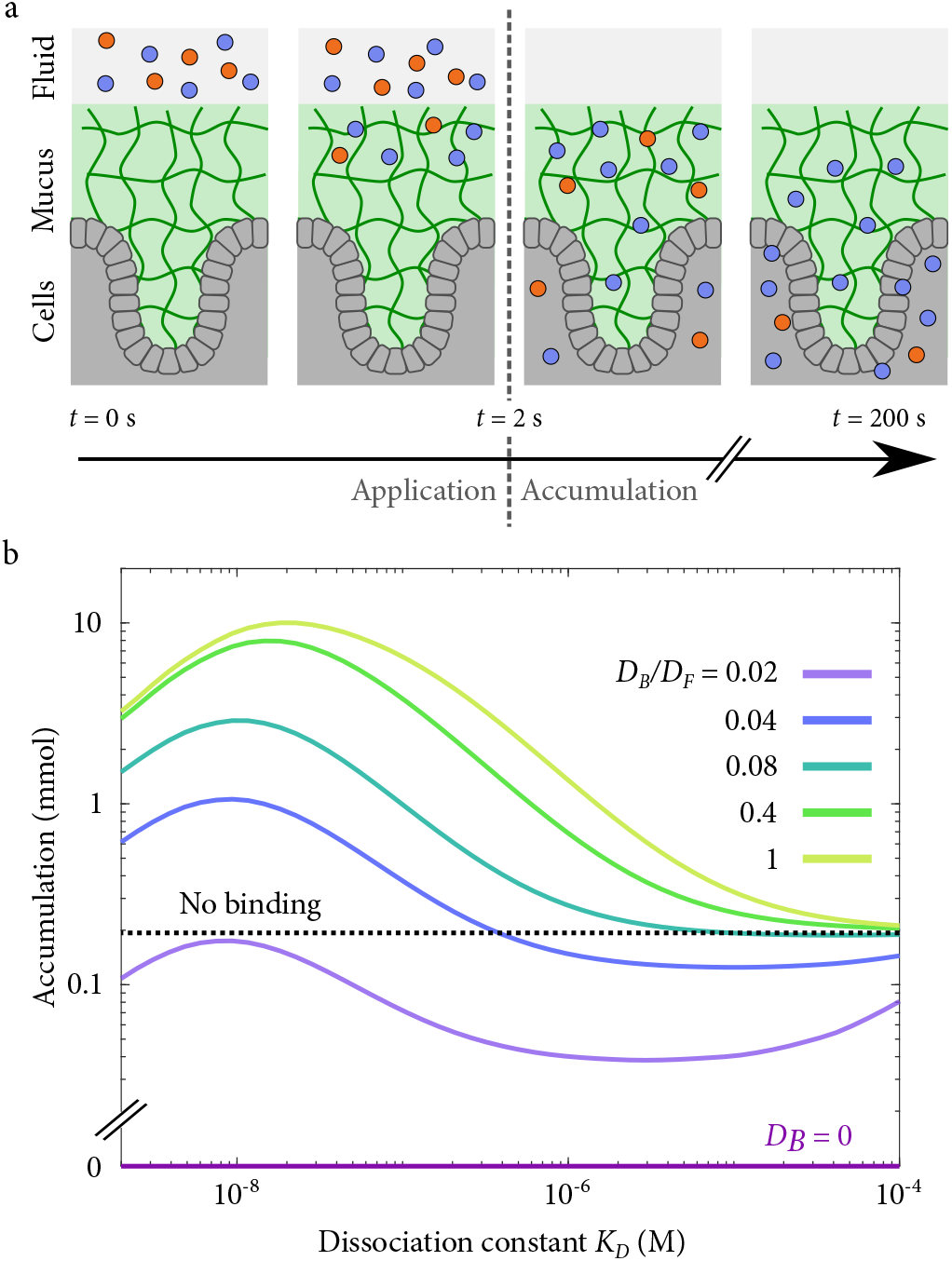
(a) Schematic of simulated transient drug application showing binding (blue) and inert (red) particles introduced to the fluid above a mucus layer (green) and subsequently washed away. Some particles are retained in the mucus layer for a time, and some of those enter the cells below (grey). Figure adapted from [67]. (b) Accumulation for a 2-s application followed by a 200-s accumulation period.

When the external reservoir is only transiently high in concentration, binding which immobilizes particles is particularly ineffective at transporting the drug to the cells. All particles that bind will have initial transient behavior with binding occurring quickly. The particles are thus delayed in reaching the tissue surface. Once the surrounding liquid is replaced, the steady loss of drug to the surrounding media is faster than diffusion to the tissue surface. For binding without diffusion, the transport to the tissue was typically orders of magnitude slower than that of an inert particle for reasonable parameter values (Fig. 5).

Remarkably, bound-state diffusion can deliver more molecules to the tissue surface than can the absence of binding (Fig. 5). The mucus binds particles and their passage to the tissue continues when the inert particles have diffused away. As with nuclear pore transport, there is an optimal binding affinity for transport. For our parameters, the optimal dissociation constant is *K_D_* ≈ 100 nM, giving a selectivity *S* ≈ 50. If possible experimentally, this increase would be similar to or greater than improvements from making particles more inert [67]. Bound diffusion would be particularly valuable in cases where the kinetics of absorption to the surface can be limiting, for example, if the concentration of cell surface receptors or internalization of bound species is low relative to the applied concentration. We do not explicitly model the rate of absorption at the tissue surface in our model. However, for the perfectly absorbing boundary conditions we consider, the instantaneous concentration at the tissue surface is steadier with binding than without (Fig. 6). Thus, the mucus itself can act as a sustained drug release platform.

**Figure 6.**
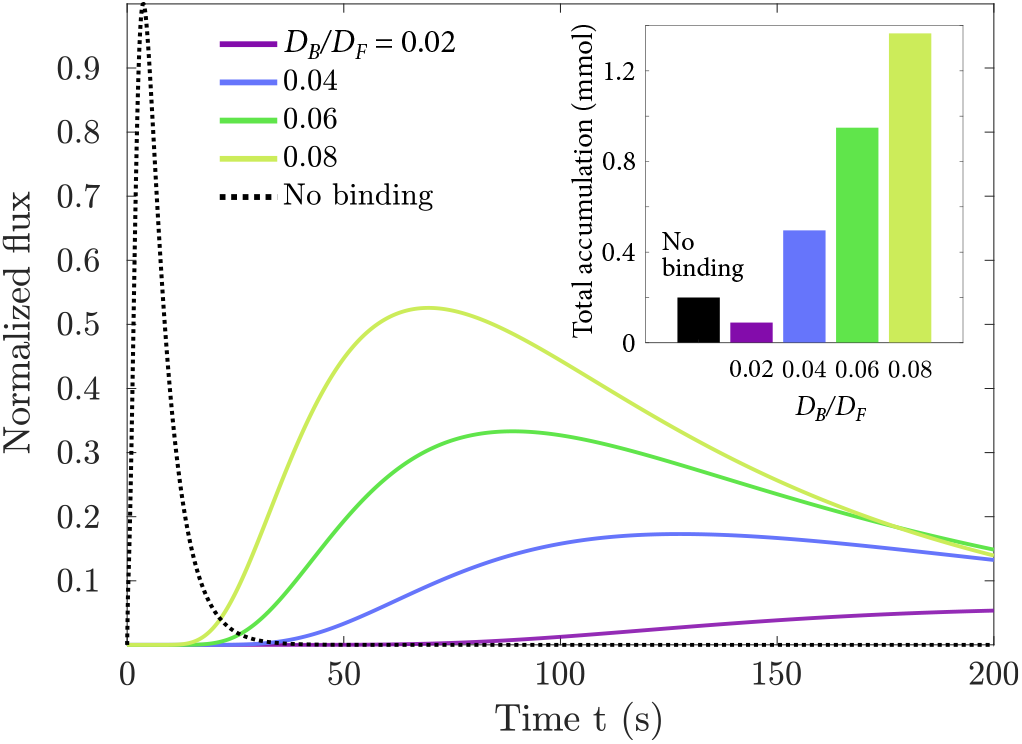
Normalized flux for a 2-s application followed by a 200-s accumulation period, shown for several values of bound diffusion constant ratio *D_B_*/*D_F_*. Inset: Total accumulated flux after 200 s as a function of *D_B_*/*D_F_*.

### Mucosal dynamics

A significant problem in drug delivery to many mucosal systems is the continuous production and sloughing of mucus. We consider an approximate model including outward flux to represent mucus sloughing: mucus and surrounding fluid are produced on one surface and disappear on the other, ignoring spatial and temporal inhomogeneities such as those occurring during coughing [69]. We considered the same transient drug application as above. No increase in transport of mucus occurs with outward flow relative to the case without outward flow, but we analyze whether bound diffusion still increases flux across the barrier even in the presence of an outward flow.

Bound diffusion can improve delivery efficacy to the underlying tissues even with outward flow. The relevant timescale for particles to diffuse across a mucus barrier of length *l_m_* is 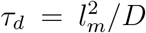, whereas the mucus is completely replaced on a timescale of *τ_r_* = *l_m_*/*v*. If *τ_d_* ≪ *τ_r_*, then bound diffusion is effective for increasing delivery through the barrier, whereas in the opposite limit it is not, as particles are removed from the barrier more rapidly than they can diffuse through it (Fig. 7). Using the parameters described in the Supplemental Material [24], bound diffusion should be effective for *v* ≪ 0.2 *μ*m/s, in agreement with the accumulation curves in Fig. 7. We estimate an outward mucus velocity of *v* ~ 10^−3^ μm/s for the eye [24], well within the regime in which bound diffusion can improve accumulation.

**Figure 7.**
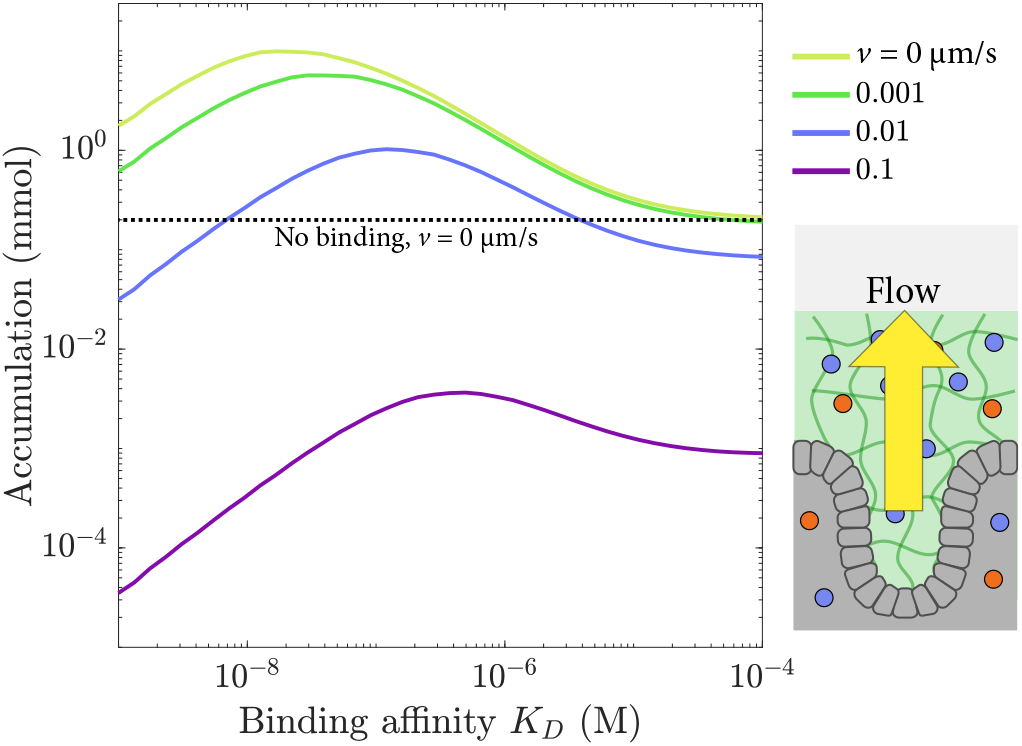
Total accumulation for a 2-s application followed by a 200-s accumulation period as a function of dissociation constant *K_D_*, shown for several values of the flow velocity v away from the barrier.

## CONCLUSIONS

In this paper we developed a hierarchical model of bound state diffusion and used physically realistic parameters and boundary conditions to predict selective transport in the NPC and mucus. Our results show that both tethered diffusion and inter-chain transfer can lead to significant selectivity in the NPC. As tethered diffusion could be engineered into nanoparticle-based drug delivery vessels, we additionally tested whether bound-diffusion based selectivity was robust to mucus-specific features such as transient application and active shedding.

We focus on dynamic multivalent binding to flexible tethers, two conserved features of the NPC that may be present in a wide variety of biological systems (DNA binding proteins, mucus, extracellular matrix, membrane-less organelles) and could be engineered into drug delivery systems [70–73]. Membrane-less organelles contain multivalent binding and flexible polymers [74], making selective transport a likely, though potentially under-appreciated, phenomenon in those systems. Our work is consistent with previous work showing that bound-state diffusion is necessary for selective transport (Fig. 2) and complements other proposals for mechanisms of bound-diffusion that include solute binding to mobile carriers [18, 20], and a hand-off between multivalent binding sites [19, 21].

We quantitatively compared the relative contributions of two mechanisms of bound diffusion: (1) tethered diffusion during transient binding events to flexible polymers and (2) inter-chain transfer of multivalent transport factors between flexible polymers. We built on significant previous interest in inter-chain transfer [21, 22] by determining its quantitative contribution to selective transport. Our model predicts that the dominant molecular mechanism of bound diffusion in the nuclear pore depends on the extent to which the FG Nups are crosslinked (Fig. 4). If the FG Nups form polymer brushes with long effective contour length (~40 nm), tethered diffusion is the primary mechanism, and is sufficient to provide high selectivity (Fig. 3). FG Nups are most likely somewhat entangled or crosslinked [43, 49, 62], which increases the importance of inter-chain hopping. However, even for tether length as short as the typical inter-FG spacing along each FG Nup (~4 nm), tethered diffusion still has the potential to contribute significantly (Fig. 3), because the fast kinetics of FG Nup-transport factor interactions mean that even motion while bound to short polymer segments can contribute to selectivity. This effect persists even if the transport factor is nearly always bound, as long as it unbinds and binds frequently; a transport factor that is bound >99% of the time to 4-nm tethers can still have selectivity from only tethered diffusion of *S* ~ 40 (Fig. 3).

Experimental measurements of NPC selectivity are quantitatively reproduced by our model of tethered diffusion, which relies on the flexibility of FG Nups and the rapid kinetics of FG Nup-transport factor binding. For conservative parameter choices, tethered diffusion produces values of the bound diffusion constant which result in selectivity of up to 200. A range of selectivity has been measured experimentally for NTF2 which agrees well with the model’s predictions (Table I), despite the minimal nature of the model.

Our model suggests that the principle of bound diffusion could be used design nanoparticles which pass more effectively through mucus barriers and provide a more uniform flux to the cells’ surface than those that are inert, potentially improving drug delivery. In our model of transient drug application, particles with bound-state mobility pass through the barrier up to 50-fold more effectively than those that do not bind, and orders of magnitude more effectively than particles that are immobilized by binding (Fig. 5). Even when the bound diffusion constant is small relative to the free diffusion constant (*D_B_* ~ *D_F_*/20), bound diffusion can increase total accumulation in the underlying tissue while also providing a more uniform flux over time, acting as a sustained-release mechanism (Fig. 6). In contrast, current nanoparticle design focuses on inert mucus-penetrating particles for drug delivery [4–6]. The addition of an outward flux of mucus to model clearance decreases but does not remove the benefits of bound diffusion for moderate values of outward velocity (Fig. 7). Nanoparticles designed for bound-state mobility within mucus barriers may therefore be able to improve drug delivery.

We thank Noel Clark, Matthew Glaser, and Franck Vernerey for useful discussions, computational resources of the BioFrontiers Institute, and funding from NIH R35 GM119755 and K25GM110486, NSF DMR1725065 and DMR1420736, the Boettcher Foundation, and a CU innovative seed grant.

