## Supplemental Material for "Design principles of selective transport through biopolymer barriers"

(Dated: July 19, 2019)

### ENTRY AND EXIT TIMES AS RATING-LIMITING STEPS

Here we analyze the model in the limit that the rate-limiting steps to transport are the entry and exit from the pore. The selectivity $S$, the ratio of the flux of the two species, is inversely proportional to the ratio of the transport time $\tau_p$. As in the work of Bickel and Bruinsma [1], we assume that entry into the pore has a free energy cost $\beta \Delta F = \nu - \epsilon$, $\nu$ is the effective free energy cost of placing an otherwise identical but non-binding particle into the pore, and $\epsilon$ is the free energy gain due to binding to the pore. Without loss of generality, we let $k_{\rm up} = k_o e^{-\beta \Delta F}$ be the rate of hopping over the barrier into the pore, and $k_{\rm down} = k_o$ be the rate of hopping out of the pore. Here $k_o$ is an attempt frequency to move the characteristic transition distance $\delta$ between the solution and the pore: $k_o = D/\delta^2$.

The total transport time for the free energy barrier is then $\tau_p = 1/k_{\rm up} + 1/k_{\rm down} = (1 + e^{\beta |\Delta F|})/k_o$, giving a selectivity

$$S = \frac{1 + e^{\nu}}{1 + e^{|\nu - \epsilon|}}. \tag{S1}$$

The selectivity is maximized when $|\Delta F| = 0$, or when the binding energy directly offsets the free energy cost of an equivalent non-binding species entering the gel. In that case, the maximum selectivity is $S_{\rm max} = (1 + e^{\nu})/2$. Additionally, if there is active release of the transport factor or cargo from the pore, the maximum selectivity remains the same, but $\epsilon$ is replaced by an effective energy gain at the exit $\epsilon^*$, in general less than $\epsilon$ if transport is to be enhanced.

### NUMERICAL INTEGRATION OF DIFFERENTIAL EQUATIONS

We numerically integrate the time-dependent reaction-diffusion equations (1, 2) using Crank-Nicolson stepping for the linear portion and forward Euler for the non-linear portion:

$$a_{n+1} = a_n + \delta t L_{op}(a_n + a_{n+1})/2 + W_{op}(a_n)\delta t, \tag{S2}$$

where $L_{op}$ is the linear component of the differential equations and $W_{op}(a)$ a non-linear function of $a$ such that $\frac{\partial a}{\partial t} = L_{op} a + W_{op}(a)$. The numerical integration was implemented in MATLAB. We numerically found steady state solutions of the equations using the differential equation solver bvp4c in MATLAB.

### FLUX IN THE LINEAR BINDING APPROXIMATION

The analytical solution for flux can be directly derived in the linear case. For ease of calculation, we reverse the concentration gradient found in the main text, so that $T(0) = 0$ and $T(L) = T_L$, allowing us to calculate flux at $x = 0$. The reaction-diffusion equations (1, 2) at steady state in the linear limit $N \approx N_t$ are

$$0 = -k_{\text{on}} N_t T + k_{\text{off}} C + D_F \frac{\partial^2 T}{\partial x^2}, \tag{S3}$$

$$0 = k_{\text{on}} N_t T - k_{\text{off}} C + D_B \frac{\partial^2 C}{\partial x^2}. \tag{S4}$$

The change of variables $C = C_x + N_t K_A T$ ($K_A = k_{\text{on}}/k_{\text{off}} = 1/K_D$ ) yields

$$0 = k_{\text{off}} C_x + D_F \frac{\partial^2 T}{\partial x^2} \tag{S5}$$

$$0 = -k_{\text{off}} C_x + D_B \frac{\partial^2 C_x}{\partial x^2} + N_t K_A D_B \frac{\partial^2 T}{\partial x^2}. \tag{S6}$$

Substituting $C_x(x) = -\frac{D_F}{k_{\text{off}}} \partial^2 T/\partial x^2$ makes equation (S6) a fourth-order ODE

$$\lambda^2 \frac{\partial^2 T}{\partial x^2} = \frac{\partial^4 T}{\partial x^4}, \tag{S7}$$

where $\lambda^2 = k_{\text{off}}(D_F + N_t K_A D_B)/(D_F D_B)$. Solutions to this equation have the form $T(x) = b + mx + fe^{\lambda x} + ge^{-\lambda x}$, where $b,\ m,\ f$ and $g$ are constants fixed by four boundary conditions: free TF concentration is fixed at the edges of the pore, with $T(0) = 0$, $T(L) = T_L$. No flux of bound TF into or out of the pore occurs, giving $\partial C/\partial x|_{x=0} = 0$, $\partial C/\partial x|_{x=L} = 0$. The constants of integration are

$$b = -(f + g), \tag{S8}$$

$$m = \frac{T_L \lambda \left(\zeta - (D_F/k_{\text{off}})\lambda^2\right)\left(e^{L\lambda} + 1\right)}{2\zeta - 2\zeta e^{L\lambda} + L\zeta\lambda - (D_F/k_{\text{off}})L\lambda^3 - (D_F/k_{\text{off}})L\lambda^3 e^{L\lambda} + L\zeta\lambda e^{L\lambda}}, \tag{S9}$$

$$f = -\frac{\zeta m}{\lambda \left(\zeta - (D_F/k_{\text{off}})\lambda^2\right)\left(e^{L\lambda} + 1\right)}, \tag{S10}$$

$$g = \frac{\zeta m + f\zeta\lambda - (D_F/k_{\text{off}}) f \lambda^3}{\zeta\lambda - (D_F/k_{\text{off}})\lambda^3}. \tag{S11}$$

where $\zeta = N_t/K_D$. This leads to a concentration profile of bound TFs

$$C(x) = \zeta \left(b + mx + fe^{\lambda x} + ge^{-\lambda x}\right) - (D_F \lambda^2/k_{\text{off}})\left(ge^{-\lambda x} + fe^{\lambda x}\right). \tag{S12}$$

To determine the selectivity, we calculate the steady-state flux out of the pore $J = -D_F \partial T/\partial x|_{x=0}$, giving

$$J = -D_F(m + \lambda f - \lambda g) \tag{S13}$$

$$J = \frac{T_L(D_F^2/k_{\text{off}})\lambda^3\left(e^{L\lambda}+1\right)}{2\zeta - 2\zeta e^{L\lambda} + L\zeta\lambda - (D_F/k_{\text{off}})L\lambda^3 - (D_F/k_{\text{off}})L\lambda^3 e^{L\lambda} + L\zeta\lambda e^{L\lambda}} \tag{S14}$$

For a non-binding particle, $C(x) = 0$, $T = T_L x/L$, and

$$J_n = -\frac{D_F T_L}{L}. \tag{S15}$$

We note that the selectivity $J/J_n$ is then independent of $T_L$ in the linear approximation.

### PARTITIONING AND THE OIL-WATER MEMBRANE

For comparisons of solubility between oil and water, the partition coefficient $\gamma$ is a ratio: the concentration of the species in oil divided by the concentration of the species in water, at equilibrium. Hydrophobic molecules (those with a high solubility in oil) have $\gamma > 1$ and are more concentrated in the oil than in the water. Hydrophilic molecules (those with a high solubility in water) have $\gamma < 1$ and are more concentrated in the water than in the oil.

In transport through the oil-water membrane, the partition coefficient also determines the flux: hydrophobic molecules have higher flux than hydrophilic. To see this in the model, note that the concentration at position $x$ within the membrane is given by $c(x)$, with $c(0) = \gamma c_L$ and $c(L) = \gamma c_R$. The flux across the membrane is given by $J = -D\frac{\partial c}{\partial x} = \gamma D(c_L - c_R)/L$, if $D$ is the diffusion coefficient within the membrane (Fig. S2).

### NPC PARAMETERS

We used the FG-filled pore length $L = 100$ nm [2, 3]; $L = 30$ nm gave similar results (data not shown). Total FG Nup concentration was determined from an estimate of the number of TF binding sites (800), and the volume of a cylinder of diameter 60 nm and length $L$, $N_t = 4.7$ mM. The free diffusion coefficient of a TF within the pore was determined from the flux of a non-binding species [4] to be $D_F = 0.12\ \mu\text{m}^2/\text{s}$. This is a reasonable value, with slower diffusion than that of karyopherins in the nuclear compartment, $D_F = 1\ \mu\text{m}^2/\text{s}$ [5]. The barrier imposed by the FG Nups to a TF and its non-binding counterpart is incorporated into the effective TF concentration at the edge of the gel. We estimate this barrier $\approx 1.5\ k_B T$ for an NTF2 sized molecule [6]. We estimate the cytoplasmic concentration of NTF2 is 5 $\mu$M. Then $T_L = 5 \times e^{-1.5}\mu\text{M} = 1\ \mu\text{M}$. The binding affinity of TFs for FG Nups within the NPC is not well known, and involves a complex interplay between site-specific affinity and change in polymer conformations [7]. Therefore, we consider a range that includes most measurements and estimates of $K_D$ between 10 nM and 10 $\mu$M [6–10].

### MUCUS BARRIER PARAMETERS

The mucus parameters were based on those experimentally measured for the eye. We calculated flux and accumulation for a mucus barrier with thickness $L = 5\ \mu$m [11]. The free diffusion constant was $D_F = 0.5\ \mu\text{m}^2/\text{s}$, corresponding roughly to a 300 nm particle diffusing in water. The concentration gradient was applied for 2 s, a reasonable time for application of drug to the eye, *e.g.* with eyedrops. The concentration gradient was then removed and the system modeled for a further 200 s. We assumed a 10 mM concentration of binding sites. The expected value of mucus velocity away from the eye was calculated using a tear secretion rate of 4 $\mu$L/min [12] and assuming an eye surface area of 1 cm$^2$, resulting in a velocity of 0.1 $\mu$m/s. As tears contain 0.01% mucins [13] and are therefore less viscous than mucus layers, which typically contain 2-5% mucins [14], we scaled the velocity proportionately by the ratio $0.01/5 = 0.002$, for a final expected velocity of $v = 2 \times 10^{-3}\ \mu\text{m/s}$.

### DIFFUSION NOTE

Here, we make the approximation that the diffusion is normal, with $\langle x(t)^2 \rangle \propto t$. In this model, motion is slightly subdiffusive because of the nonlinearity of the equations, and because binding to a random location within a well introduces a fluctuating force. However, in the regimes most relevant to the NPC, where the binding lifetime is short, this effect is minimal.

### HOPPING SIMULATION

In our simulation of TF motion with hopping between FG Nups while bound, we represented each FG Nup as an entropic spring (represented by a harmonic potential well). Well positions were randomly chosen from a uniform distribution, with the exception that we always placed one well at the starting position of the TF. The particle (the TF) started the simulation bound to this FG Nup, and remained bound to a Nup throughout the simulation. While bound to one FG Nup, the TF diffused within the harmonic well representing that FG Nup. We recorded the position and mean-squared displacement of the TF from its starting location, which we then used to determine a bound diffusion coefficient, as described in more detail below. The TF could hop between tethers by changing the well in which it moved.

#### Diffusion in a potential well

The TF moved in the harmonic potential of the FG Nup according to Brownian dynamics. TF position updated using a force-dependent diffusive step [15, 16]

$$x(t+\delta t) = x(t) + \frac{F}{\Gamma}\delta t + \delta x, \tag{S16}$$

where $F$ is the force acting on the particle, $\Gamma$ is the drag coefficient, $\delta t$ is the timestep, and $\delta x$ is a random Brownian step drawn from a Gaussian distribution with variance $\sigma^2 = 2D\delta t$. The drag coefficient of a spherical particle at low Reynolds number is given by Stokes' Law as $\Gamma = 6\pi\eta r$, where $\eta$ is the fluid's viscosity and $r$ is the sphere's radius. This result can be combined with the Einstein relation $D = k_BT/(6\pi\eta r)$ to give

$$\Gamma = \frac{k_BT}{D}. \tag{S17}$$

The force $F = -k\Delta x$, where $k$ is the spring constant of the FG Nup and $\Delta x$ is the displacement of the particle from the Nup attachment point. We model the FG Nup as a worm-like-chain at small extension, so that $k = 3k_BT/(2\ell_pL_c)$, where $\ell_p$ is the tether persistence length and $L_c$ is the contour length. Then

$$x(t+\delta t) = x(t) - \frac{3D\Delta x\delta t}{2\ell_pL_c} + \delta x = x(t) - DK\Delta x\delta t + \delta x, \tag{S18}$$

where $K$ is the normalized spring constant $K = k/k_BT = 3/(2\ell_pL_c)$.

#### Hopping probability

We constructed the hopping probability $P_{\text{hop}}$ to satisfy the principle of detailed balance. During every iteration of the simulation, we picked an FG Nup at random from a list of the $M$ Nups near enough to have a reasonable probability of hopping. TF hopping to the new FG Nup was attempted with success probability

$$P_{\text{hop}} = r_{\text{hop}} M \delta t e^{-\Delta G/2}. \tag{S19}$$

Here the base hopping rate $r_{\text{hop}}$ is a dimensionless input parameter, and the change in free energy (in units of $k_B T$) between the current Nup and the proposed new Nup is

$$\Delta G = \frac{1}{2} K (x - x_{\text{new}})^2 - \frac{1}{2} K (x - x_{\text{cur}})^2, \tag{S20}$$

where $K$ is the normalized spring constant, $x$ is the particle's current position, $x_{\text{cur}}$ is the anchor location of the Nup to which the particle is currently bound, and $x_{\text{new}}$ is the anchor location of the proposed new Nup. Note that when a hop succeeded, the energy landscape changed to that of the new Nup, but the TF's position did not change during the hop. There is no upper bound on $P_{\text{hop}}$, but we adjusted the timestep to ensure that $P_{\text{hop}}$ was greater than unity no more than 0.5% of the time that a hop was attempted.

#### Mean-squared displacement and diffusion coefficient calculation

We ran each simulation for $10^7$ time steps with $\delta t = 0.01$ $\mu$s and recorded the particle's position every 100 time steps. We calculated the mean-squared displacement (MSD) $\langle x^2 \rangle$ of the TF and averaged it over 100 runs [Fig. S7(a)]. We then computed

$$\rho_{\text{MSD}}(t) = \langle x^2(t) \rangle \rho(k_{\text{off}}, t) = k_{\text{off}} \langle x^2(t) \rangle e^{-k_{\text{off}} t}, \tag{S21}$$

as shown in Fig. S7(b), and numerically integrated the distribution in time. We determined the bound diffusion coefficient from the typical MSD-per-binding-event $\overline{\langle x^2 \rangle}$ using

$$D_B = \frac{k_{\text{off}} \overline{\langle x^2 \rangle}}{2}. \tag{S22}$$

Here, the factor of 1/2 is appropriate because we consider a one-dimensional random walk.

| Nup fragment | Nup concentration | Molecule | MW | Partition coeff. | Diffusion coeff. in gel | Diffusion coeff. free | $K_D$ | $D_B$ | $D_B/D_F$ | $S$ | $S$ with 10x partition coeff. | Notes |
|---|---|---|---|---|---|---|---|---|---|---|---|---|
| Nsp1 (2-601) | 3 mM | IBB-MBP-mEGFP-ImpB<br>MBP-mCherry | 510 kD<br>70 kD | 100<br>0.16 | 0.17 $\mu$m$^2$/s | 4.03 $\mu$m$^2$/s | 4.8 $\mu$M | 0.17 $\mu$m$^2$/s | 0.04 | 42 | 50 | [17] |
| Nup57 (1-223)–Nup49 (1-246) | 3.7 mM | IBB-MBP-mEGFP-ImpB<br>MBP-mCherry | 510 kD<br>70 kD | 400<br>0.15 | 0.1 $\mu$m$^2$/s | 2.7 $\mu$m$^2$/s | 1.4 $\mu$M | 0.1 $\mu$m$^2$/s | 0.04 | 69 | 45 | [17] |
| Nup57 (1-223) –Nsp1 (2-601)–Nup49 (1-246) | 1.7 mM | IBB-MBP-mEGFP-ImpB<br>MBP-mCherry | 510 kD<br>70 kD | 350<br>0.1 | 0.24 $\mu$m$^2$/s | 4.03$\mu$m$^2$/s | 0.48 $\mu$M | 0.24 $\mu$m$^2$/s | 0.06 | 38 | 16 | [17] |
| Nsp1 (2-175) | 3.0 mM | IBB-MBP-mEGFP-ImpB<br>MBP-mCherry | 510 kD<br>70 kD | 100<br>3 | 0.04 $\mu$m$^2$/s | 12.1$\mu$m$^2$/s | 90 $\mu$M | 0.04 $\mu$m$^2$/s | 0.003 | 1.4 | 4.3 | [18] |
| Nsp1 (2-601) | 3.0 mM | IBB-MBP-mEGFP-ImpB<br>MBP-mCherry | 510 kD<br>70 kD | 60<br>0.4 | 0.22 $\mu$m$^2$/s | 6.94$\mu$m$^2$/s | 20 $\mu$M | 0.22 $\mu$m$^2$/s | 0.03 | 15 | 40 | [18] |
| Nsp1 (1-601) | 2.2 mM | IBB-Redstar-ImpB<br>IBB-Redstar | 530 kD<br>150 kD | 1000<br>0.3 | 0.1 $\mu$m$^2$/s | 0.2$\mu$m$^2$/s | 0.66 $\mu$M | 0.1 $\mu$m$^2$/s | 0.5 | 230 | 100 | [19] |
| Nsp1 (1-601) | 2.2 mM | GFP-ImpB<br>IBB-Redstar | 124 kD<br>150 kD | 100<br>0.3 | 0.1-0.2 $\mu$m$^2$/s | 0.2$\mu$m$^2$/s | 6.6 $\mu$M | 0.1-0.2 $\mu$m$^2$/s | 0.5-1 | 210-250 | 230-240 | [19] |
| Nsp1 (1-601) | 2.2 mM | GFP-ImpB<br>acRedStar | 124 kD<br>117 kD | 100<br>0.05 | 0.1-0.2 $\mu$m$^2$/s | 0.2-1$\mu$m$^2$/s | 1.1 $\mu$M | 0.1-0.2 $\mu$m$^2$/s | 0.1-1 | 94-260 | 53-130 | [19] |
| P - Nsp1 (274-601) - P * | 4.4 mM | IBB-MBP-mEGFP-ImpB<br>IBB-MBP-mEGFP | 510 kD<br>100 kD | 7<br>0.9 | 2.78 $\mu$m$^2$/s | 16.0 $\mu$m$^2$/s | 560 $\mu$M | 2.42 $\mu$m$^2$/s | 0.15 | 5.3 | 25 | [20] |

TABLE S1. Predicted selectivity $S$ (for a 100-nm barrier) of FG Nup hydrogels in previous work. We took partition and diffusion coefficients from tables in references or calculated them using concentration plots. We determined the dissociation constant $K_D$ from the partition coefficient of the binding species ($P_B$) and non-binding species ($P_N$) and the Nup concentration $N_t$ using $K_D \approx (P_I/P_B)N_t$. Note that the measured $P_B$ is an underestimate of the true partition coefficient. We estimated the bound diffusion coefficient from the in-gel (effective) diffusion coefficient ($D_{\text{eff}}$) and the probability of the binding species being bound ($p_b \approx 1 - K_D/N_t$) using $D_B = p_b D_{\text{eff}}$. We used the reaction-diffusion equations discussed in the main text to estimate the selectivity. Because partition coefficient estimates were lower bounds, we also calculated selectivity assuming that the reported partition coefficients were 10% of their actual value. P - Nsp1 (274-601) - P * refers to a fusion between Nsp1 (274-601) and a pentameric coiled-coil P which facilitates the aggregation of the Nsp1 domain into hydrogels. See [20].

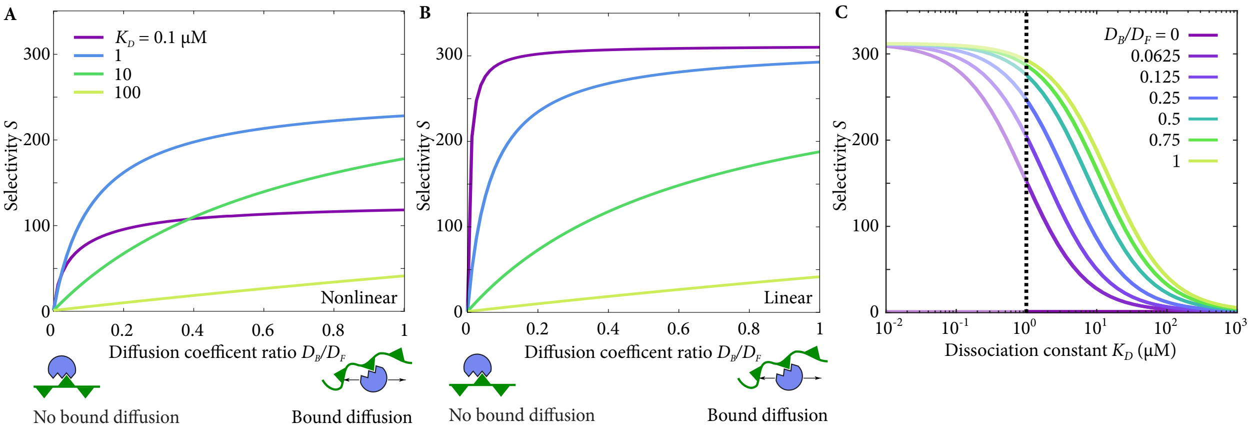


FIG. S1. (a, b) Selectivity as a function of diffusion coefficient ratio, with varying dissociation constant, in the full nonlinear model (a) or in the linear approximation (b). (c) Selectivity as a function of dissociation constant, with varying diffusion coefficient ratio, in the linear approximation. The region to the left of the dotted line is non-physical; the full nonlinear solution should be used.

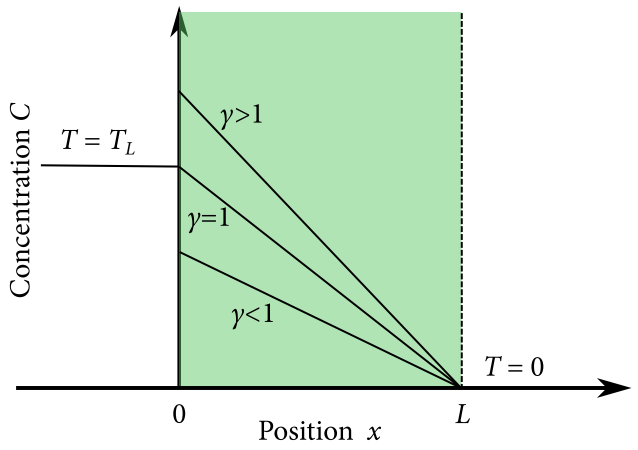


FIG. S2. Concentration $C$ vs position $x$ across an oil membrane of length $L$ surrounded by water. Several values of the partition coefficient $\gamma$ are shown.

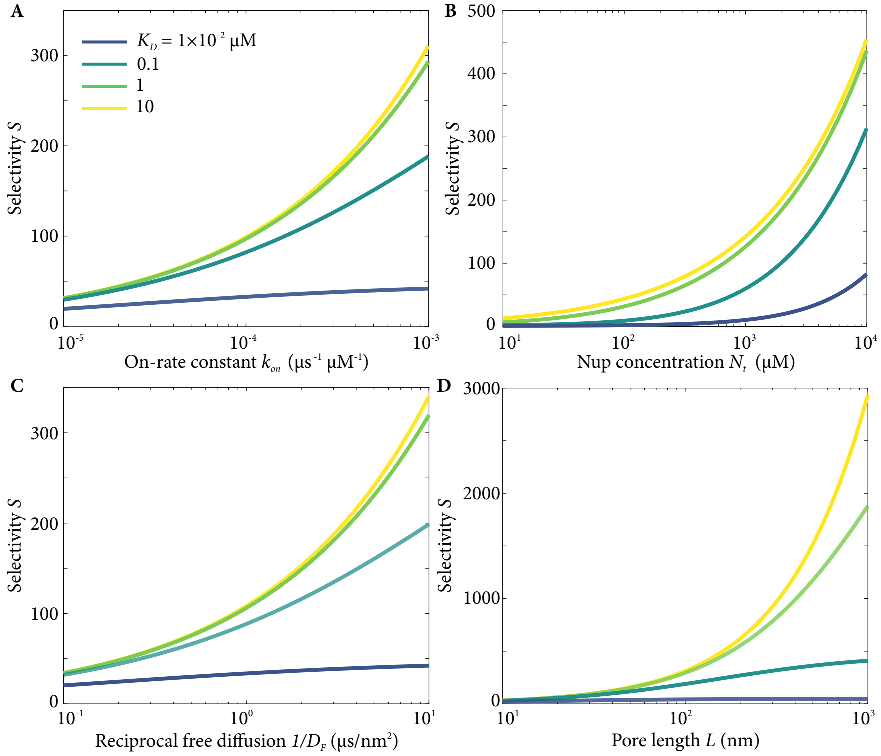


FIG. S3. Dependence of selectivity on variation of individual parameters: (a) on-rate constant, (b) total FT Nup concentration, (c) inverse of the free diffusion coefficient, and (d) pore length, with varying dissociation constant. All values calculated using the linear solution to the binding-diffusion equations. Bound diffusion coefficient $D_B = 0.1 D_F$. Other parameters fixed at values from the NPC parameters section.

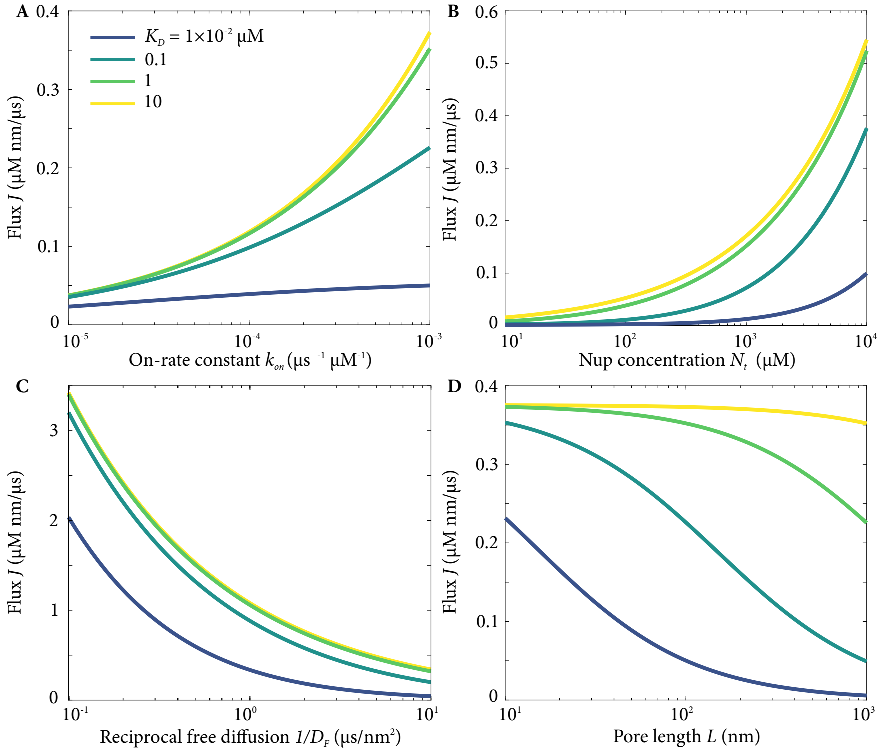


FIG. S4. Dependence of flux on variation of individual parameters: (a) on-rate constant, (b) total FT Nup concentration $N_t$, (c) inverse of the free diffusion coefficient, and (d) pore length, with varying dissociation constant. All values calculated using the linear solution to the binding-diffusion equations. Bound diffusion coefficient $D_B = 0.1D_F$. Other parameters fixed at values from the NPC parameters section.

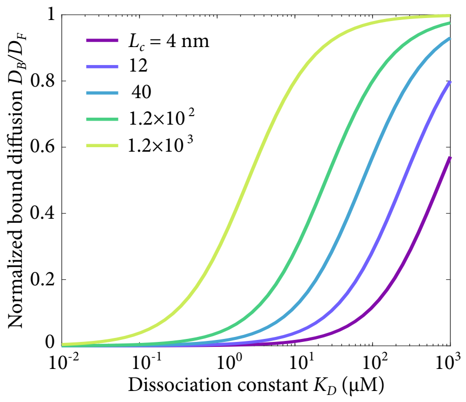


FIG. S5. Bound diffusion coefficient $D_B$ vs dissociation constant $K_D$ using the tethered-diffusion model. Several values of the Nup contour length $L_c$ are shown.

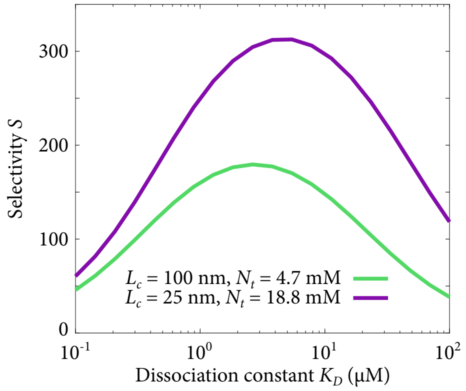


FIG. S6. Selectivity as a function of dissociation constant in the tethered diffusion model, varying Nup contour length $L_c$ and total Nup concentration $N_t$. The product $L_c N_t$ is held constant.

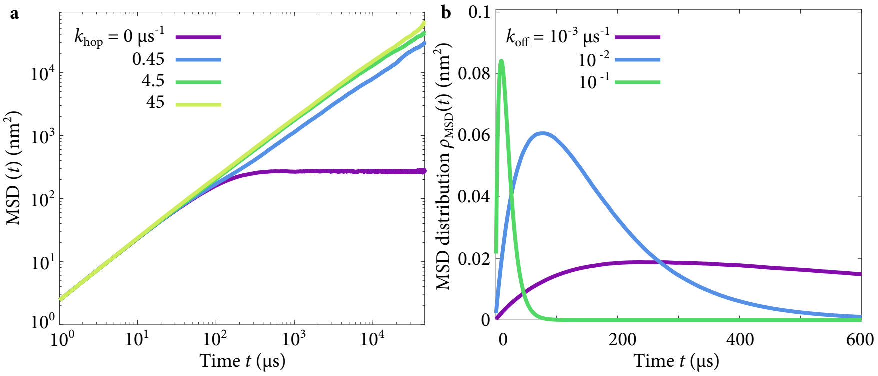


FIG. S7. (a) Examples of mean-squared displacement (MSD) of a simulated TF in the inter-chain hopping model, with varying hopping rate. (b) Examples of MSD distributions $\rho_{\text{MSD}}(t)$ used in estimating the diffusion coefficient, with varying unbinding rate. Tethers have 40 nm contour length; other parameters are as discussed in the main text.

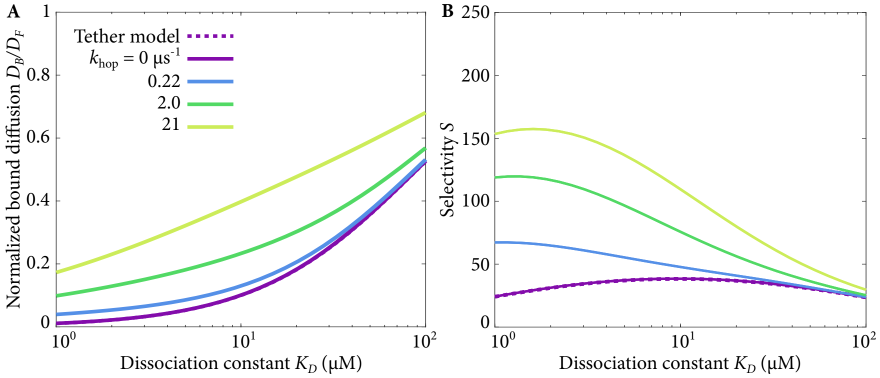


FIG. S8. Bound diffusion and selectivity as a function of dissociation constant, with varying hopping rate for FG Nups with $L_c = 4$ nm.

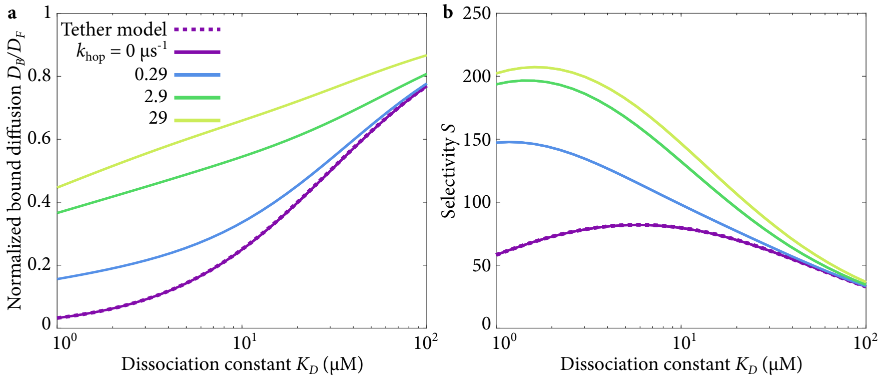


FIG. S9. (a) Bound diffusion and (b) selectivity as a function of dissociation constant, with varying hopping rate for FG Nups with $L_c = 12$ nm.

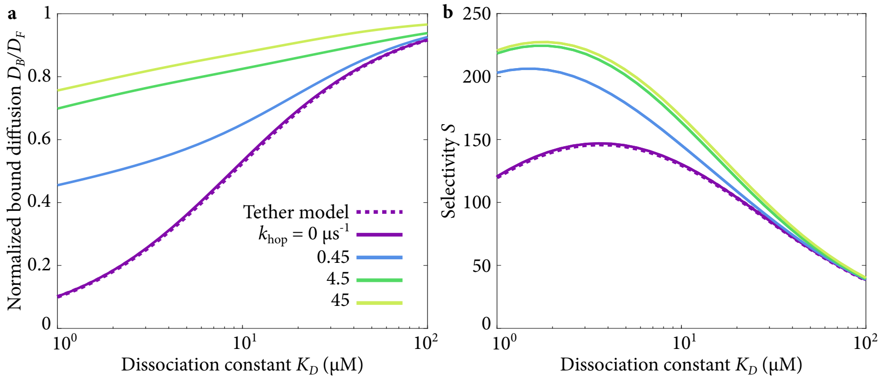


FIG. S10. (a) Bound diffusion and (b) selectivity as a function of dissociation constant, with varying hopping rate for FG Nups with $L_c = 40$ nm.

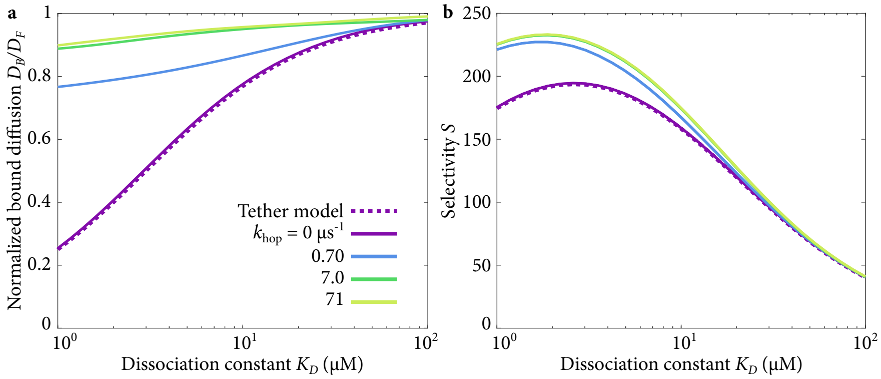


FIG. S11. (a) Bound diffusion and (b) selectivity as a function of dissociation constant, with varying hopping rate for FG Nups with $L_c = 120$ nm.

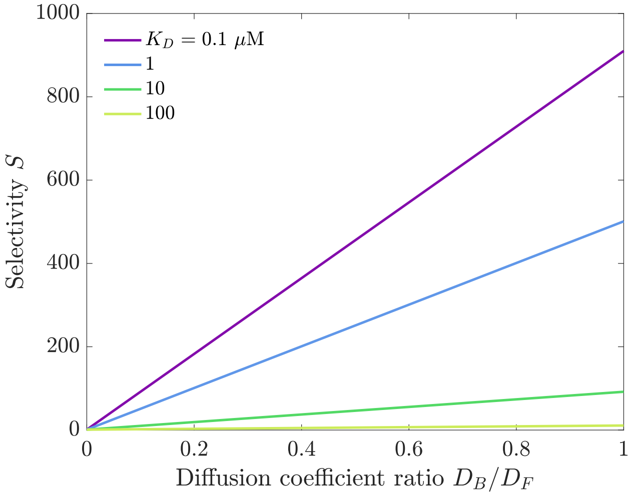


FIG. S12. Selectivity $S$ as a function of diffusion coefficient ratio $D_B/D_F$ using the model described in Yang *et al.* [21]. The transport factor concentration in the reservoir is $T_L = 1\ \mu$M and the total Nup concentration is $N_t = 1$ mM.
